# Effects of date fruit (*Phoenix dactylifera*) on sperm cell morphology and reproductive hormonal profiles in cypermethrin-induced male infertility

**DOI:** 10.1101/2020.06.17.156687

**Authors:** Ubah Simon Azubuike, Agbonu Oluwa Adikpe, Columbus Philemon Kwinjoh, Abah Kenneth Owoicho, Chibuogwu Ijeoma Chika, Abalaka Samson Eneojo, Abayomi Samuel Bankole, Enem Simon Ikechukwu, Ajayi Itopa Etudaye

## Abstract

Date fruits are endowed with medicinal values, including boosting the male fertility status, but with meagre empirical evidence. Thus, the current study was designed to assess the ameliorative and potential adverse effects of date fruit extracts (*Phoenix dactylifera*) on cypermethrin-induced male infertility. The study was conducted in two phases using adult male Wistar rats (n = 42, 180 – 220 g and aged 14 - 16 weeks). The first phase was a single oral dose toxicity study to ascertain the suitability of date fruit extract and cypermethrin administered at 250 mg/kg and 60 mg/kg, respectively. The second phase, which included four treatment groups of six animals per group, assessed the effects of date fruits on cypermethrin-induced infertility. At the termination of the experiment, semen was collected by epididymal extraction for the assessment of sperm abnormalities, motility, mass activity, semen pH, and percentage live. Serum samples were also collected for testosterone and follicle stimulating hormone (FSH) profiling, and the collected data was subjected to statistical analysis. The group administered only cypermethrin showed a decrease in percentage motility, live, mass activity and an increase in total abnormalities over the control group while the group exposed to only date fruits extracts showed increased percentage motility, live, mass activity and a decrease in total abnormalities over the control. The results of a combined administration of date fruit extracts and cypermethrin on a separate group showed a consistently reduced percentage of anatomically abnormal sperm cells and a general improvement of sperm motility and mass activity. There was no significant difference in the weight of the Wister rats in all the groups (p > 0.05). However, testosterone and FSH levels were significantly reduced (p < 0.05) by date fruit extract treatment. The current report provides evidence of the potential ameliorative effects of date fruit extracts in cypermethrin-induced male infertility and cautions excessive use or abuse since some adverse effects were observed.

## 1.0. Introduction

Cypermethrin is a neurotoxic synthetic pyrethroid that is commonly used as a synergist to increase the potency of insecticides on pests [1]. Sequel to a recommendation by the World Health Organisation (WHO), the use of the chemical has become popular for the effective treatment of mosquitoes in Asia and Africa [2], where malaria is epidemic [3]. However, studies have reported cases of acquired infertility following exposure of cypermethrin to animals [4 – 8]. The lethal dose (LD_50_) of cypermethrin in rats following oral administration is 251 mg/kg of body weight [9], but periodic subacute doses over prolonged periods result in chronic genotoxic effects including a significant reduction in the relative weights of the testes, epididymis, seminal vesicle and prostate glands that, consequently, disrupts the normal androgen status and androgenesis [10 – 13]. Thus, irrespective of the value of cypermethrin in agriculture, the residues, in trace amounts or higher levels, in the air, soil and water, are toxic to mammals.

However, there are indications that date fruits, *Phoenix dactylifera*, possesses some ameliorative effects over cypermethrin-induced infertility, but these theories have not been scientifically validated. *Phoenix dactylifera* is a monocotyledon that belongs to the family of Arecaceae. The plant and its fruits hold high economic values in the tropical and desert areas of Africa, the Middle East and Asia [14], and are endowed with several medicinal values [15]. For instance, it enhances the relief of oxidative stress by neutralizing free radicals and decomposing peroxidases [16]. Interestingly, studies have demonstrated a strong correlation between cypermethrin exposure and the production of excess reactive oxygen species (ROS). An excessive production of ROS is a known cause of oxidative stress [17] that induces organ injury by the oxidative damage of deoxyribonucleic acid (DNA), proteins and lipids [18]. Juxtaposing these reports with the antioxidant properties of *Phoenix dactylifera* suggests that date fruits could possess an ameliorative influence over the pathophysiologic effects generated through chronic exposure to cypermethrin. Furthermore, date fruit extracts contain estrogenic materials such as gonad-stimulating compounds that enhance male fertility [19 – 23]. This information partly validates the tradition of eating date fruits as a fresh vegetable to boost male fertility. Together, these factors bear significance in traditional African medicine and culture, and so preparations of date fruits are often administered as oral suspensions to cure male infertility with varying degrees of successes. However, in the literature, there is scanty empirical evidence of the protective effects of date fruit on sperm parameters. Moreover, being an exogenous source of steroid with a capacity to up-regulate spermatogenesis [24], the extracts of date fruits may also disrupt normal hormonal processes. Yet, there is no evidence of the potential adverse effects of *Phoenix dactylifera* on the male reproductive system.

Therefore, the current study aimed to evaluate the therapeutic and potential adverse effects of date fruit extract on semen picture, FSH and testosterone profiles following cypermethrin-induced toxicity.

## 2.0. Materials and methods

### 2.1. Experimental animals and study area

The University of Abuja Institutional Animal Ethics Committee, Nigeria, approved the experiments conducted in the current study. The care and use of animals were also guided by the standard principles of laboratory animal care. Male Wister rats (n = 42; 180 – 220 g and aged 14 - 16 weeks) were used for the study. The animals were acclamatised to the study environment for 3 weeks prior to the commencement of the experiments. Standard plastic cages were used to house the animals and these were placed under 12:12 hour light: dark cycle. Food and water were provided *ad libitum*.

### 2.2. Plant identification, extraction and standardization

Date fruits were obtained from Shuwari, Dutse LGA, Jigawa State, Nigeria (11°42^′^04^″^N 9°20^′^31^″^E). The plant was identified at the Herbarium Centre of the University of Abuja, Nigeria, with herbarium number UNIBUJA/H/70. Debris was separated from the selected date fruits and then the fruits were allowed to air dry for a period of two weeks. The seeds were then removed from the pods and macerated using a laboratory blender (Conair^TM^ 7011S). One kilogram of the sample was weighed using a weighing balance and then poured into a 3-Litre beaker containing 2.5 L of methanol. The preparation was then left to stand at room temperature for 72 hours after which a muscling cloth was used to filter out the extract from the shaft. The extract was concentrated on a water bath until dried. The percentage yield was calculated and recorded as 35 %. A phytochemical analysis of the extract was done using a UV visible double beam spectrophotometer (Cecil 750, Cambridge England^®^).

### 2.3. Experimental design

The experiments in the current study were divided into two phases. The first phase was a single oral dose toxicity study while the second phase examined the effects of date fruit extracts on cypermethrin-induced male infertility.

#### 2.3.1. Single oral dose toxicity study

This phase of the experiment was conducted to ascertain the suitability of the selected doses of cypermethrin and date fruits to be used in the current experiment. The dose of cypermethrin used in this phase of the study was selected as a fraction of the oral LD_50_ dose for rats [9]. Following a two-week period of acclimatisation, Wister rats (n = 18) were divided into three groups of 6 animals each designated as A, B and C. Group A served as control and was administered distilled water. The test rats (Groups B and C) were fasted for 4 hours with free access to water only. Group B was administered date fruit extract orally at a dosage of 250 mg/kg of body weight and Group C was administered cypermethrin at a dosage of 60 mg/kg of body weight. Rats were observed for signs of acute toxicity including, behavioral changes and mortality for 14 days.

#### 2.3.2. Effects of date fruit extracts on cypermethrin-induced male infertility

The experimental setup for this phase comprised of four treatment groups of six animals each: Animals in Group 1 (The control group) were administered distilled water. Animals in Group 2 were administered cypermethrin at a dose of 60 mg/kg. Animals in Group 3 were administered a combination of cypermethrin and date fruit extracts at the dose rates of 60 mg/kg and 250 mg/kg, respectively. In Group 4, animals were administered date fruit extract at a dose of 250 mg/kg. All the preparations were orally administered weekly for a period of eight weeks using the Gavage method [25]. At the termination of the experiment, the rats were euthanised and semen was collected.

### 2.4. Semen collection and evaluation

Semen collection was done by epididymal extraction [26] for the assessment of sperm motility, mass activity, semen pH, percentage live and abnormalities. Serum samples were also collected for testosterone and FSH assay. A descriptive summary of epididymal extraction, according to Turner and Giles, 1982 [26] is as follows: The rats were euthanised with an over dose of anesthetic ether (Nandkrishna Chemicals). The testes were then exteriorized by incising the scrotum, and the testicles were extracted as the *tunica vaginalis* was opened. The caudal epididymis was then isolated and cleaned with warm normal saline at 37 °C, and punctured with a sterile needle to ‘milk-out’ semen. The semen was evaluated according to a method described by Zemjanis [27]. Briefly, a drop of semen was extracted onto a pre-warm glass slide. An approximately equal drop of normal saline was immediately added to dilute the semen. The semen sample was then cover-slipped and viewed using a field microscope at x4 magnification to determine the mass activity and x10 magnification to assess the progressive motility of each sample. Morphological abnormalities were determined by diluting a drop of semen with 4 % buffered formal saline stained with Eosin-Nigrosin stain. The slides were allowed to dry and then viewed under a light microscope to assess 100 sperm cells per animal using oil immersion for percentage live and abnormalities. Further, a drop of raw undiluted semen sample was taken and placed on a litmus paper for pH determination.

Photomicrographs were obtained with a light microscope (Olympus BX 53) equipped with an Olympus DP72 camera. Digital images were minimally optimised by adjusting contrast and brightness using Adobe Photoshop CC (version 2017.1.1). Vector-based illustrations were made using Adobe Illustrator CC (version 2017.1.1)

### 2.5. Serum collection and hormone assay

Hormonal assay was done using the serum sample obtained from the rats. Blood samples were collected in non-heparinized tubes through the ocular media cantus after adequately anesthetizing the rats. Clear serum samples were then collected by allowing the blood to clot and centrifuging at 1500 rpm (Anke^®^TGL-12B). The collected serum samples were stored in bottles at -20 oC until the time for analysis. Testosterone and FSH were analysed using a standard immunoassay technique, Spectra Testosterone and FSH kits (Orion Diagnostica; Finland and DRG Instruments GmbH; Germany).

### 2.6. Statistical analysis

All the data obtained were expressed as mean ± standard error of mean (SEM). The data was analyzed using one-way analysis of variance (ANOVA) and values of p < 0.05 were considered statistically significant.

## 3.0. Results

### 3.1. Single oral dose toxicity study

There was no observable sign of toxicity and mortality in groups A (distilled water) and B (date fruit extracts). However, in group C (Cypermethrin), the rats displayed signs of mild toxicity each time cypermethrin was orally administered. The signs of toxicity included jumping, uncoordinated movement and restlessness within the cage, which subsided after approximately 10 minutes.

### 3.2. Phytochemical analysis

Phytochemical analysis indicated that date fruit extract contains 18.2 % alkaloids, 19.2 % tannins, 14.6 % flavonoid, 17.8 % phenol and negligible quantities of glucoside.

### 3.3. Observed anatomic defects of sperm cells

The morphology of sperm cells in all the groups were assessed and the percentages of normal and abnormal cells were derived from a total of 2, 000 cells. Normal sperm cells were identified by the possession of a smooth hook-shaped head and a droplet-like structure located at the mid-piece of a sperm flagellum. Normal cells were observed in 76.4 % of the sperm cells present in the control samples (Figure 1d) while the abnormal cells constituted 23.6 %. However, in the group treated with cypermethrin alone, the total number of abnormal sperm cells increased by 2.2 % while groups 3 and 4 had a significant reduction in the percentage of abnormal sperm cells (Figure 1d). Further, based on the location of the abnormality, sperm cell abnormalities were grouped into head, mid-piece and tail abnormalities. The predominant head abnormality was detached heads, which reflected as detached hook-shaped structures without the other components (mid-piece and tail) of the sperm cells. The mid-piece and tail abnormalities observed included bent mid-pieces and tails, double cytoplasmic droplets, coiled tails, dag effects and free tails (Figures 2 and 3).

**Figure 1.**
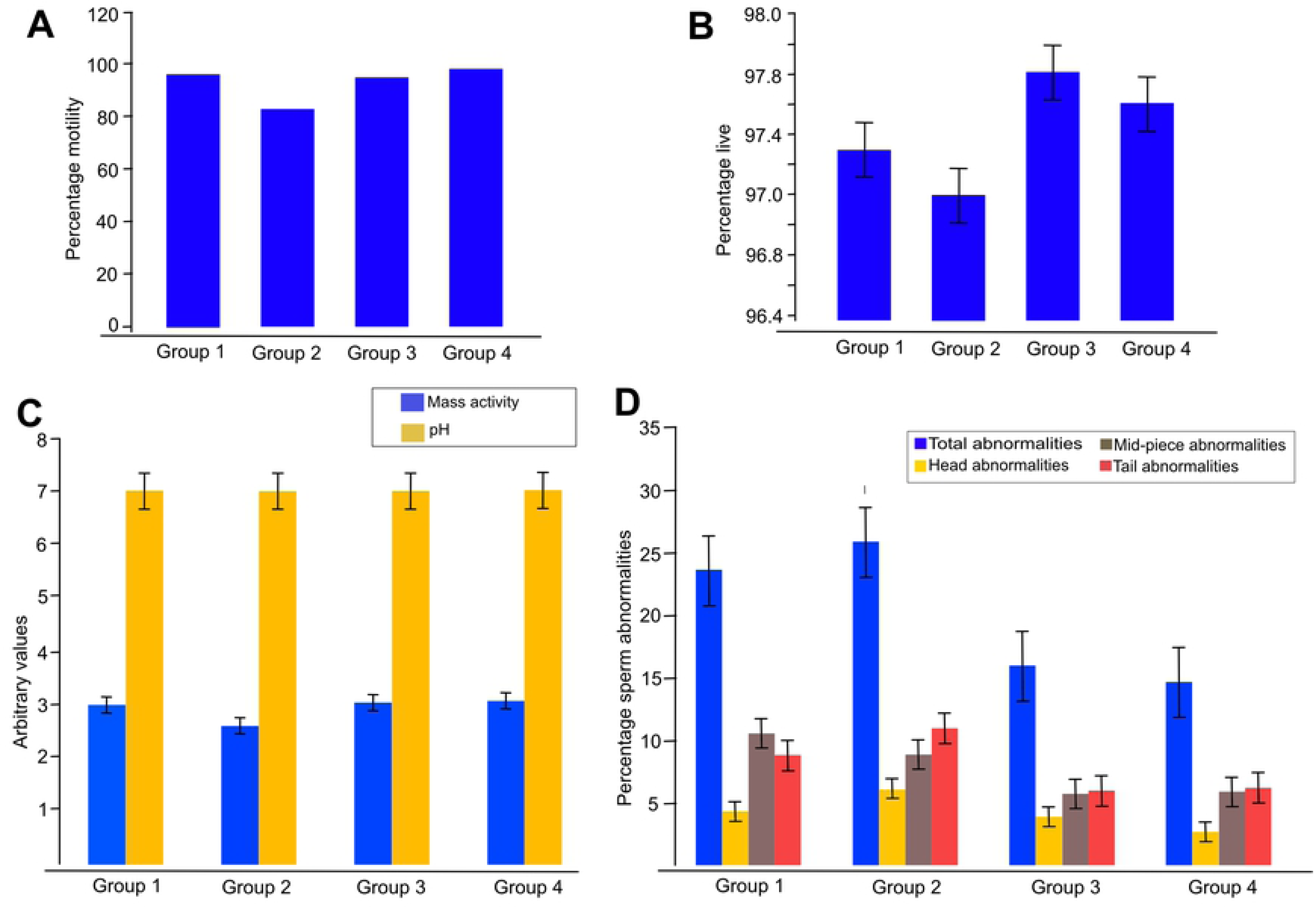
Analysis of the effects of date fruits (*Phoenix dactylifera)* on cypermethrin-induced male infertility (n = 24). The parameters of infertility measured in the current study include percentage motility (**A**), Percentage live (**B**), Mass activity and pH, and percentage sperm cell abnormalities (**D**). The results suggest a significant level of cytotoxic activity by cypermethrin on the male reproductive system as indicated in group 2 of all the parameters measured with the exception of pH. However, there is also a significant ameliorative effect following the administration of date fruit extracts as indicated in groups 3 and 4 of all the parameters measured.

**Figure 2.**
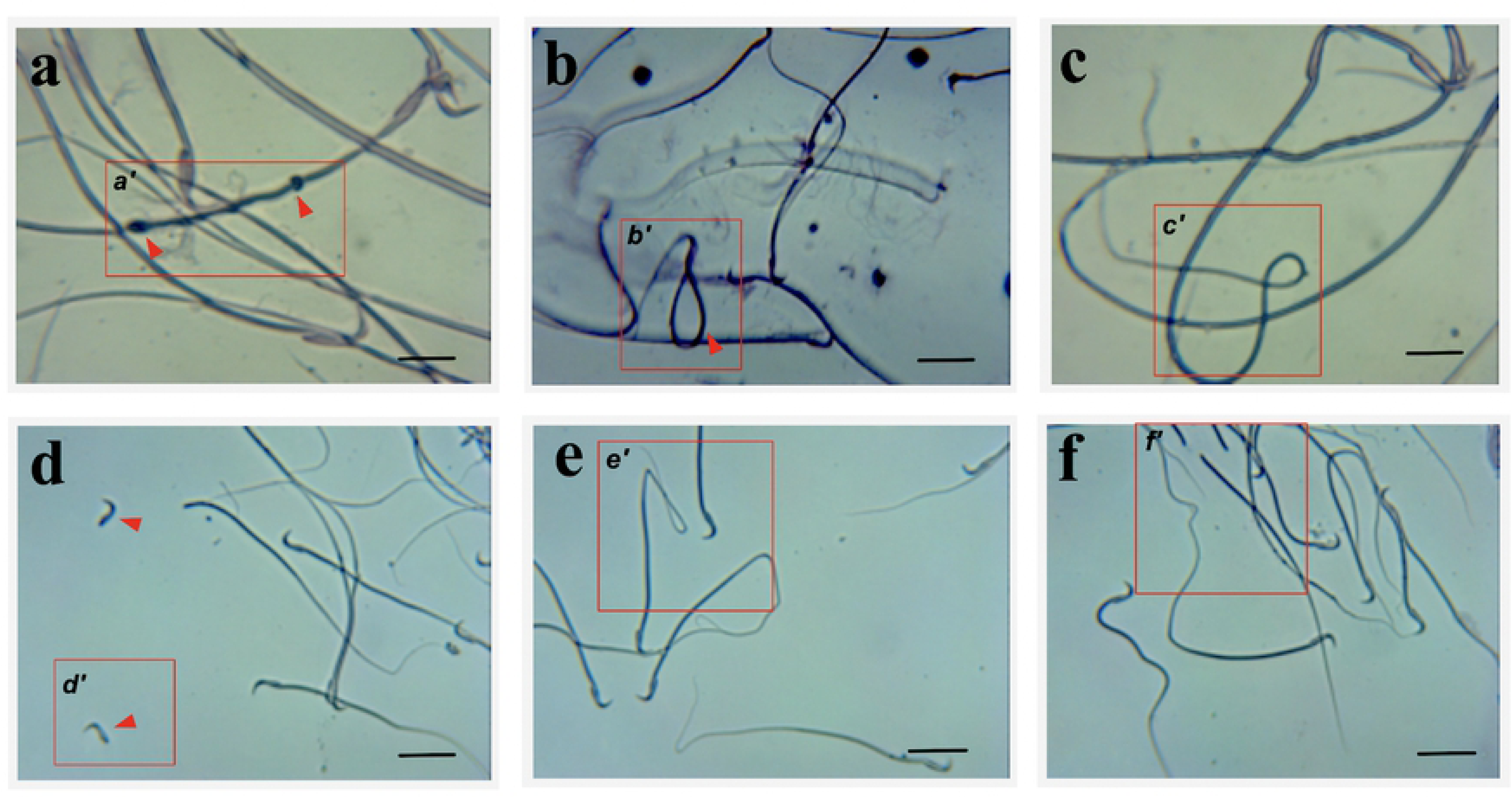
Morphological abnormalities of sperm cells in cypermethrin-induced male infertility. The pictures are representative photomicrographs of the observed sperm cell defects in the current study with highlighted areas that contain the sperm cell structural abnormality (red box and arrows). The observed abnormalities include (a) double cytoplasmic droplet, (b) Dag effect, (c) coiled tail, (d) detached head, (e) bent tails and (f) free tail. The assessment of the structural abnormalities was done on an overall reference population of 2,000 sperm cells. Scale bar = 20 µm.

**Figure 3.**
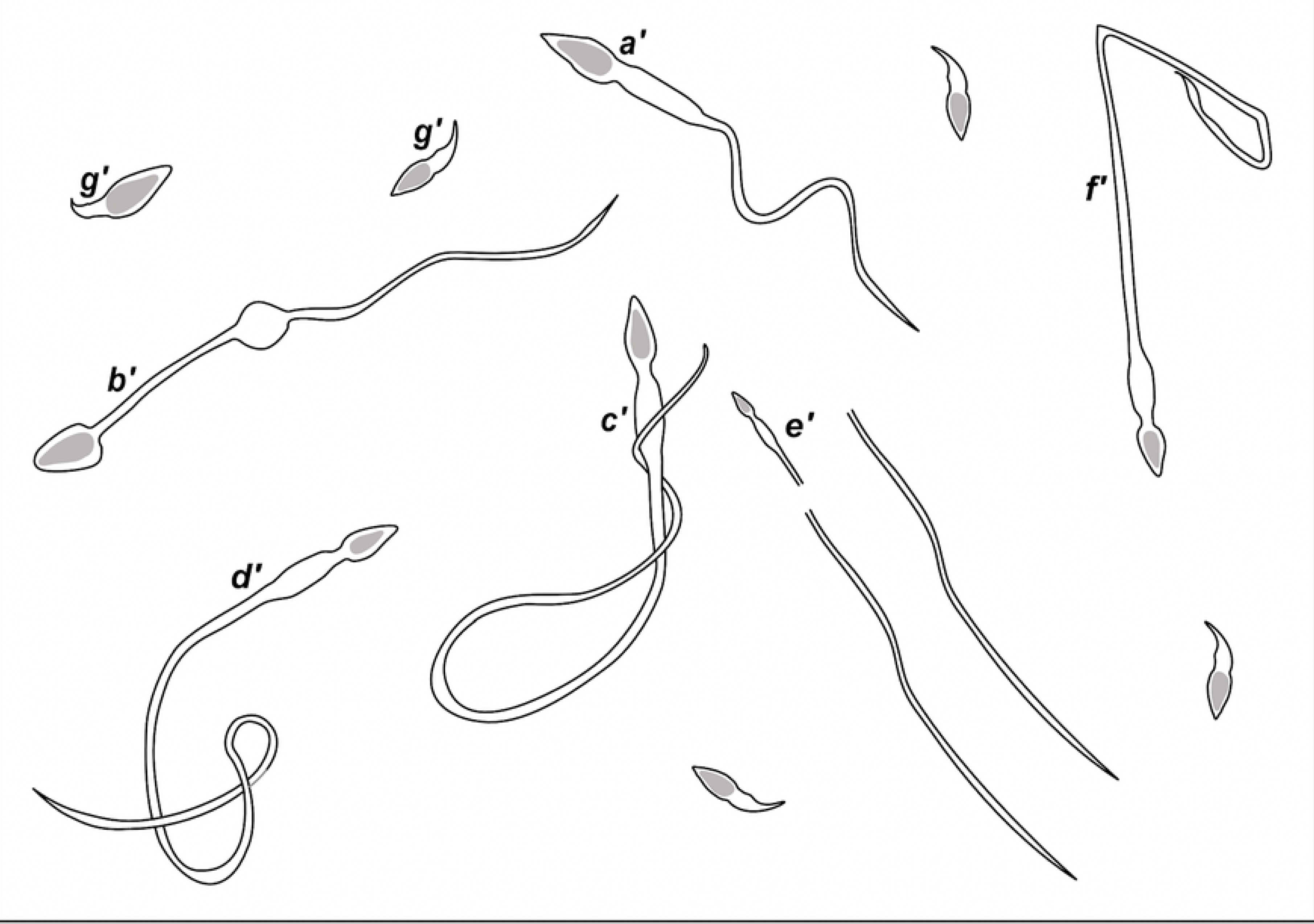
Graphical illustrations of a normal sperm cell (a) and all the structural sperm cell abnormalities (*b’ – d’*) observed in the current study. Structural abnormalities include double cytoplasmic droplets (***b’***), Dag effect (***c’***), coiled tails (***d’***), free tail (***e’***), bent tail and mid-pieces (***f’***), and detached head (***g’***). The varying percentages of these structural abnormalities in the treatment groups suggest that date fruits possess some ameliorative properties on cypermethrin-induced infertility in males.

### 3.4. Sperm motility and percentage liveability

Results showed that there was no significant difference in the sperm motility, percentage live, mass activity and pH for samples in all the groups (p > 0.05) (Figure 1a - c). There was also no significant difference in the weight of Wister rats (p > 0.05).

### 3.5. Hormonal assay

The hormonal assay showed significantly lower FSH values of 1.75 ± 0.97^b^ and 2.26 ± 1.05^b^ mIU/ml for groups 3 and 4, respectively (p < 0.05). Testosterone profiles also showed significantly lower values of 0.77 ± 0.09b ng/ml for group 3 (p < 0.05) (Table 1). A summary of final testosterone and FSH profile against the reference values [28] are presented in Table 1.

**Table 1.** Follicle stimulating hormone (FSH) and testosterone profiles of Wister rats (n = 20) Mean (±SEM) Body weight, FSH and Testosterone profiles of Wister rats tested for the effect of date fruit extracts on Cypermethrin induced male impotence.

| Groups | FSH (mIU/ml) * | Testosterone (ng/ml) ** |
| --- | --- | --- |
| Group 1 | $6.18 \pm 2.48^a$ | $2.00 \pm 0.51^a$ |
| Group 2 | $8.04 \pm 4.37^a$ | $1.72 \pm 0.61^a$ |
| Group 3 | $1.75 \pm 0.97^b$ | $0.77 \pm 0.09^b$ |
| Group 4 | $2.26 \pm 1.05^b$ | $1.25 \pm 0.37^a$ |

## 4.0. Discussion

In the current study, the capacity of cypermethrin to cause damage to the male reproductive cells and modulate the endocrinal system at a fraction of the median oral LD_50_ [9] in rats was validated in the group of animals that were exposed to cypermethrin alone (Figure 1a and b). This finding aligns with a substantial number of studies that have identified cypermethrin as a primary cause of infertility [16, 29, 30, 31]. The authors report that chronic exposure to cypermethrin could result in cytotoxic organ damage since a microscopic assessment of sperm cells revealed a cocktail of sperm cell morphological abnormalities (Figures 2 and 3). Although the rats administered cypermethrin exhibited signs of mild toxicity, the signs lasted a short duration after which apparent health was restored indicating that the selected dose of 60 mg/kg was suitable for a more chronic experiment. The mechanisms through which cypermethrin exerts its actions includes direct damage to the male reproductive system facilitated by oxidative stress [29, 30], disruption of neurotransmission [31] and the modulation of endocrinal balance [16]. The current study did not investigate any of these mechanisms of action since the experiments focused on the protective effects of date fruit extracts. However, the novelty herein is the observation that date fruits reduces the pathologic effects observed in cypermethrin-induced infertility. Evidence to support this statement includes the observed general improvement of spermatozoa health reflected in the higher percentage of motility and liveability, and the lower sperm cell abnormalities observed when animals treated with date fruits were compared with the control values.

Infertility is a challenging global issue [32], and the current study reveals substantial evidence of the protective values of date fruits over the condition. For instance, a reduction in the incidence of anatomically abnormal sperm cells (Figure 1d) when rats were treated with date fruit extracts is a scientific evidence indicating capacity to reduce the chronic cytotoxic damage of cypermethrin on the male reproductive system. The most common anatomic defect observed (mid-piece and tail abnormalities, which appeared either detached, inadequately developed, coiled or bent) can be attributed to weak or incomplete adhesion between the plasma membrane and nuclear envelope at the posterior ring of the mid-piece [33]. These defects are common during spermatogenesis and often occur at the implantation fossa of the mid-piece [34]. The defects are phenotypic reflections of gene interactions and mutation. Moreover, the defects indicate the degree of abnormality and immaturity of the sperm population in an ejaculate, which have a directly proportional relationship with the fertility status of an animal. Interestingly, the differences in the number of structurally defective sperm cells between the groups were consistent, and our findings agree with those of previous studies [35, 36], which reported that date palm pollen possess properties that improve male infertility.

Another valuable evidence that date fruit extracts reduces the impact of cypermethrin on the toxicity of sperm cells was indicated by the increased motility or mass activity observed in the current study. Increased motility could result from the antioxidant property of date fruit extracts against cypermethrin-induced oxidative stress since date fruit extract contains antioxidants such as coumaric and ferulic acids [37]. The authors suggest that similar antioxidant properties were responsible for the improved percentage liveability.

However, the adverse effects of date fruit extracts on the reproductive hormones was a cause for concern as it appeared that the extracts possess the capacity to modulate the endocrine system by significantly lowering FSH and testosterone hormonal levels below referenced values (Table 1). The endocrinal mechanisms that resulted in this effect are unclear, but it is known that date fruits possess estrogenic and gonadotropic activities [21, 22], which classifies the plant as a source of exogenous steroid. So, if ingested for a prolonged period, date fruits potentially increase systemic estrogen levels, thereby triggering a negative feedback mechanism of exogenous gonadotrophin at the level of the anterior pituitary. This feedback loop consequently reduces endogenous gonadotropins including FSH, Sertoli cells levels, androgen binding proteins and testosterone levels [38]. Thus, date fruits bear potential adverse effects on fertility.

### 4.1. Conclusion

The causes of infertility are diverse, and range from innate genetic disorders [39, 40] to several acquired factors including infection [41, 42], chemo cytotoxicity and endocrinal imbalance [43]. The results from the current study show that ingested date fruit extracts possess some ameliorative properties over cypermethrin-induced male infertility. Indicators of fertility such as sperm cell morphology, motility and mass activity supported the claim of an ameliorative effect. However, identifying a specific treatment regimen is difficult since the aim of an ideal strategy would require targeting the root cause of the disease, while minimizing adverse effects. Amidst the protective properties of date fruit extracts, the plant also carried some adverse effects on the fertility of the male reproductive system, as observed in the present study. The endocrinal system was modulated by a reduction in the levels of testosterone and FSH. These changes were indicative of adverse effects. Against these back-drop, it would be important for future studies to establish a dose-effect relationship. A dose-effect relationship curve is one of the criteria for determining causality, which would strengthen the inferences made in the current study. Also, investigations into the physiologic mechanisms recruited by date fruit extracts (*Phoenix dactylifera*) would be crucial to fully understand the medicinal value of the plant.

## Acknowledgement

The authors thank the staff The Sheda Science and Technology Complex (SHESTCO), Abuja, for the technical support received.

## Author contribution

All authors contributed equally to the study.

### Funding

No funding

